# Single-cell RNA expression profiling of ACE2, the receptor of SARS-CoV-2

**DOI:** 10.1101/2020.01.26.919985

**Authors:** Yu Zhao, Zixian Zhao, Yujia Wang, Yueqing Zhou, Yu Ma, Wei Zuo

## Abstract

A novel coronavirus SARS-CoV-2 was identified in Wuhan, Hubei Province, China in December of 2019. According to WHO report, this new coronavirus has resulted in 76,392 confirmed infections and 2,348 deaths in China by 22 February, 2020, with additional patients being identified in a rapidly growing number internationally. SARS-CoV-2 was reported to share the same receptor, Angiotensin-converting enzyme 2 (ACE2), with SARS-CoV. Here based on the public database and the state-of-the-art single-cell RNA-Seq technique, we analyzed the ACE2 RNA expression profile in the normal human lungs. The result indicates that the ACE2 virus receptor expression is concentrated in a small population of type II alveolar cells (AT2). Surprisingly, we found that this population of ACE2-expressing AT2 also highly expressed many other genes that positively regulating viral entry, reproduction and transmission. This study provides a biological background for the epidemic investigation of the COVID-19, and could be informative for future anti-ACE2 therapeutic strategy development.

---

SARS-CoV-2 is a coronavirus identified as the cause of an outbreak of coronavirus disease 2019 (COVID-19), which now causes death in approximately 2~3% of infected individuals in China (1–4). Patients with confirmed infection have reported respiratory illness such as fever, cough, and shortness of breath (5). Once contacted with the human airway, the spike proteins of this virus can associate with the surface receptors of sensitive cells, which mediated the entrance of the virus into target cells for further replication. Xu et.al., first modeled the spike protein to identify the receptor for SARS-CoV-2, and indicated that Angiotensin-converting enzyme 2 (ACE2) could be the receptor for this virus (6). ACE2 is previously known as the receptor for SARS-CoV and NL63 (7–9). Following studies focusing on genome sequence and structure of the receptor-binding domain of the spike proteins further confirmed that the new coronavirus can efficiently use ACE2 as a receptor for cellular entry, with estimated 10- to 20-fold higher affinity to ACE2 than SARS-CoV (10, 11). Zhou et. al. conducted virus infectivity studies and showed that ACE2 is essential for SARS-CoV-2 to enter HeLa cells (12). These data indicated that ACE2 is the receptor for SARS-CoV-2.

The tissue expression and distribution of the receptor decide the tropism of virus infection, which has a major implication for understanding the pathogenesis and designing therapeutic strategies. Previous studies have investigated the RNA expression of ACE2 in 72 human tissues and demonstrated its expression in lung and other organs (13). Lung is a complex organ with multiple types of cells, so such real-time PCR RNA profiling based on bulk tissue could mask the ACE2 expression in each type of cell in the human lung. The ACE2 protein level was also investigated by immunostaining in lung and other organs (13, 14). These studies showed that in normal human lung, ACE2 is mainly expressed by type II and type I alveolar epithelial cells. Endothelial cells were also reported to be ACE2 positive. Immunostaining detection is a reliable method for identification of protein distribution, yet accurate quantification remains a challenge for such analysis. The recently developed single-cell RNA sequencing (scRNA-Seq) technology enables us to study the ACE2 expression in each cell type and provides quantitative information at single-cell resolution. Previous work has built up the online database for scRNA-Seq analysis of 8 normal human lung transplant donors (15). In current work, we used the updated bioinformatics tools to analyze the data. Some of the results of these studies have been previously reported in the form of preprint (doi: https://doi.org/10.1101/2020.01.26.919985) (16).

We analyzed 43,134 cells derived from normal lung tissue of 8 adult donors (Figure 1A). We performed unsupervised graph-based clustering (Seurat version 2.3.4) and for each individual, we identified 8~11 transcriptionally distinct cell clusters based on their marker gene expression profile. Typically the clusters include type II alveolar cells (AT2), type I alveolar cells (AT1), airway epithelial cells (ciliated cells and Club cells), fibroblasts, endothelial cells and various types of immune cells. The cell cluster map of a representative donor (Asian male, 55-year-old) was visualized using t-distributed stochastic neighbor embedding (tSNE) as shown in Figure 1B.

**Figure 1.**
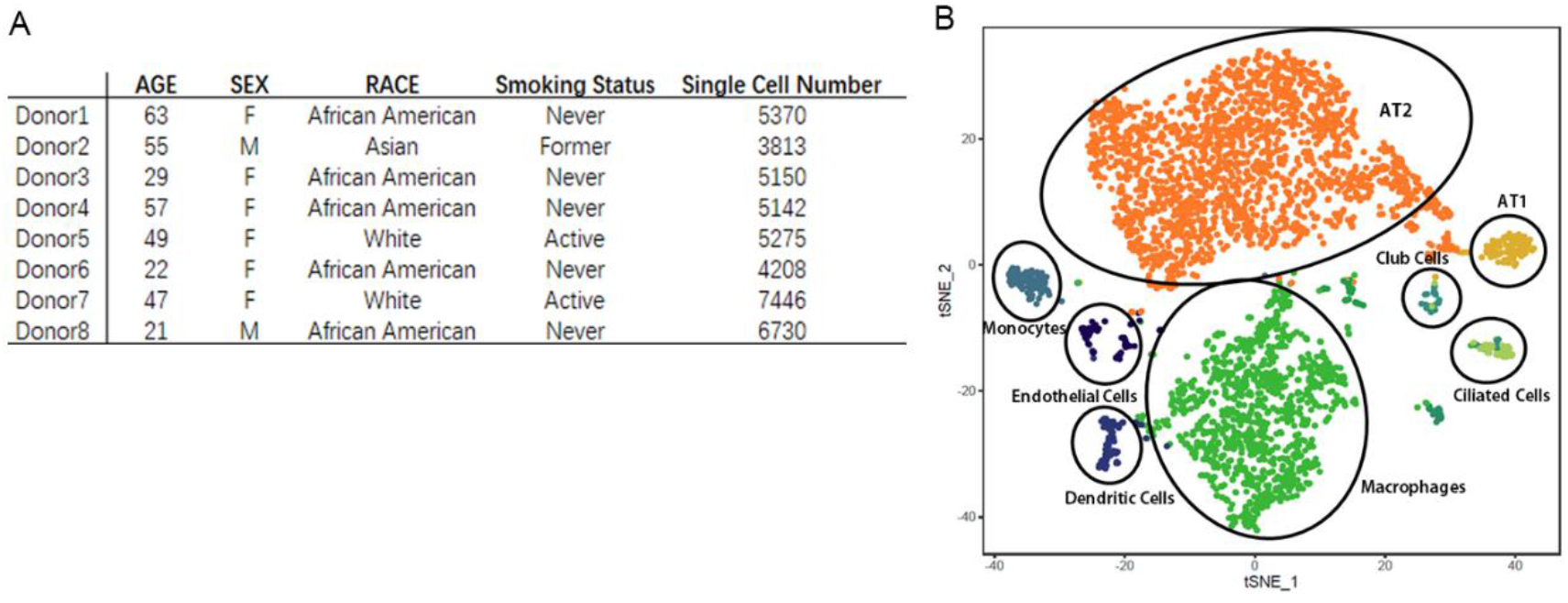
Single-cell RNA-Seq analysis of normal human lungs. (A) Characteristics of lung transplant donors for single-cell RNA-Seq analysis. (B) Cellular cluster map of the Asian male. All 8 samples were analyzed using the Seurat R package. Cells were clustered using a graph-based shared nearest neighbor clustering approach and visualized using a t-distributed Stochastic Neighbor Embedding (tSNE) plot.

Next, we analyzed the cell-type-specific expression pattern of ACE2 in each individual. For all donors, ACE2 is expressed in 0.64% of all human lung cells. The majority of the ACE2-expressing cells (83% in average) are AT2 cells. Other ACE2 expressing cells include AT1 cells, airway epithelial cells, fibroblasts, endothelial cells, and macrophages. However, their ACE2-expressing cell ratio is relatively low and variable among individuals. For the representative donor (Asian male, 55-year-old), the expressions of ACE2 and cell-type-specific markers in each cluster are demonstrated in Figure 2A.

**Figure 2.**
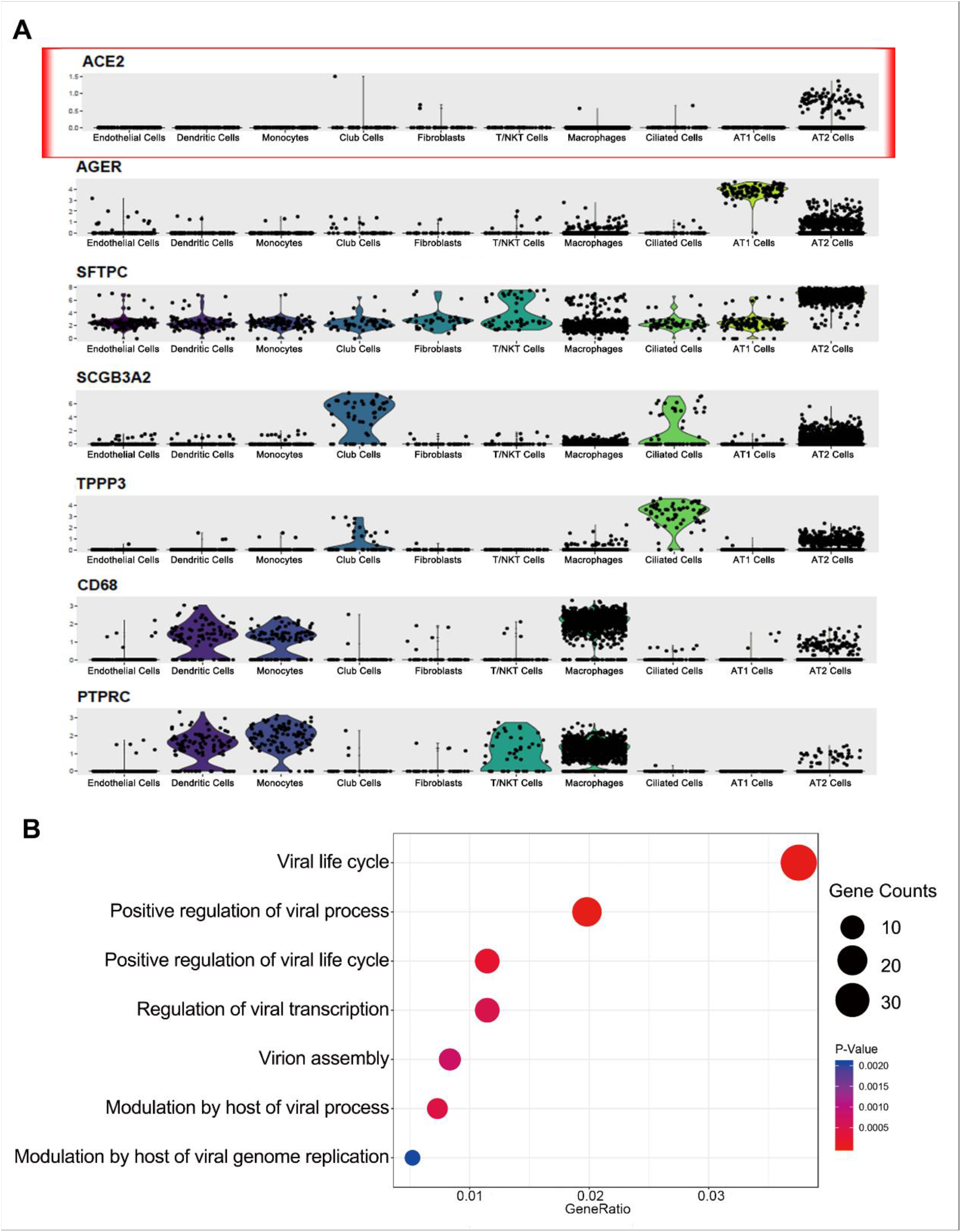
Gene expression analysis in ACE2-expressing AT2 population. (A) Violin plots of expression for ACE2 and select cell type-specific marker genes significantly upregulated in distinct lung cell clusters from an Asian male donor. AGER, type I alveolar cell marker; SFTPC (SPC), type II alveolar cell marker; SCGB3A2, Club cell marker; TPPP3, ciliated cell marker; CD68, macrophage marker; PTPRC (CD45), pan-immune cell marker. (B) Dot plot of GO enrichment analysis demonstrating enriched virus-related biological processes in the ACE2-expressing AT2 population.

There are 1.4±0.4% of AT2 cells expressing ACE2. To further understand the special population of ACE2-expressing AT2, we performed gene ontology (GO) enrichment analysis to study which biological processes are involved with this cell population by comparing them with the AT2 cells not expressing ACE2. Surprisingly, we found that multiple viral life cycle-related functions are significantly over-represented in ACE2-expressing AT2, including those relevant to viral replication and transmission (Figure 2B). We found upregulation of CAV2 and ITGB6 gene in ACE2-expressing AT2. These genes are component of lipid raft/caveolae, which is a special subcellular structure on plasma membrane critical to the internalization of various viruses including SARS-CoV (17–19). We also found enrichment of multiple ESCRT machinery gene members (including CHMP3, CHMP5, CHMP1A, VPS37B) in ACE2-expressing AT2, which were related to virus budding and release (20, 21). These data showed that this small population of ACE2-expressing AT2 is particularly prone to SARS-CoV-2 virus infection.

We further analyzed each donor and their ACE2 expressing patterns. As the sample size is very small, no significant association was detected between the ACE2-expressing cell number and any characteristics of the individual donors. But we did notice that one donor has 5-fold higher ACE2-expressing cell ratio than average. The observation on this case suggested that the ACE2-expressing profile heterogeneity might exist between individuals, which could make some individual more vulnerable to SARS-CoV-2 than others. However, these data need to be interpreted very cautiously due to the very small sample size of current dataset, and larger cohort study is necessary to draw conclusion.

Altogether, in the current study, we report the RNA expression profile of ACE2 in the human lung at single-cell resolution. Our analysis suggested that the expression of ACE2 is concentrated in a special small population of AT2 which also expresses many other genes favoring the viral infection process. It seems that SARS-CoV-2 has cleverly evolved to hijack this population of AT2 cells for its reproduction and transmission. Targeting AT2 explained the severe alveolar damage and minimal upper airway symptom after infection by SARS-CoV-2. The demonstration of the distinct number and distribution of ACE2-expressing cell population in different cohorts can potentially help to identify the susceptible population in future. The shortcoming of the study is the small sample number, and that the current technique can only analyze the RNA level but not the protein level of single cells. Furthermore, while previous studies reported abundant ACE2 expression in pulmonary endothelial cells (13, 22), we did not observe high ACE2 RNA level in this population. This inconsistency may be partly due to the fact that the cell number and portion of endothelial cells in current dataset is relatively smaller than expected. Indeed, due to the limitation of sample collection and processing, the analyzed cells in this study may not fully represent the whole lung cell population. Future quantitative analysis at the transcriptomic and proteomic level in a larger total population of cells is needed to further dissect the ACE2 expression profile, which could eventually lead to novel anti-infective strategies such as ACE2 protein blockade or ACE2-expressing cell ablation.

## Methods

Public datasets (GEO: GSE122960) were used for bioinformatics analysis. First, Seurat (version 2.3.4) was used to read a combined gene-barcode matrix of all samples. Low-quality cells with less than 200 or more than 6,000 detected genes were removed, or if their mitochondrial gene content was > 10%. Only genes found to be expressing in > 3 cells were retained. For normalization, the combined gene-barcode matrix was scaled by total unique molecular identifiers (UMI) counts, multiplied by 10,000 and transformed to log space. The highly variable genes were identified using the function FindVariableGenes. Variants arising from number of UMIs and percentage of mitochondrial genes were regressed out by specifying the vars.to.regress argument in Seurat function ScaleData. The expression level of highly variable genes in the cells was scaled and centered along each gene, and was conducted to principal component (PC) analysis.

Then the number of PCs to be included in downstream analysis was assessed by (1) plotting the cumulative standard deviations accounted for each PC using the function PCElbowPlot in Seurat to identify the ‘knee’ point at a PC number after which successive PCs explain diminishing degrees of variance, and (2) by exploring primary sources of heterogeneity in the datasets using the PC Heatmap function in Seurat. Based on these two methods, the first top significant PCs were selected for two-dimensional t-Distributed Stochastic Neighbor Embedding (tSNE), implemented by the Seurat software with the default parameters. FindClusters was used in Seurat to identify cell clusters for each sample. Following clustering and visualization with tSNE, initial clusters were subjected to inspection and merging based on the similarity of marker genes and a function for measuring phylogenetic identity using BuildClusterTree in Seurat. Identification of cell clusters was performed on the final aligned object guided by marker genes. To identify the marker genes, differential expression analysis was performed by the function FindAllMarkers in Seurat with Wilcoxon rank sum test. Differentially expressed genes that were expressed at least in 25% cells within the cluster and with a fold change more than 0.25 (log scale) were considered to be marker genes. tSNE plots and violin plots were generated using Seurat. Gene Ontology (GO) enrichment analysis of differentially expressed genes was implemented by the ClusterProfiler R package. GO terms with corrected P value less than 0.05 were considered significantly enriched by differentially expressed genes. Dot plots were used to visualize enriched terms by the enrichplot R package.

## Acknowledgements

W. Zuo designed the project; Y. Zhao, Z. Zhao, Y. Wang, Y. Zhou and Y. Ma performed the analysis; Y. Wang and W. Zuo drafted the manuscript. We thank Alexander Misharin group for sharing their original scRNA-Seq dataset to public. This work was funded by the National Key Research and Development Program of China (2017YFA0104600 to W. Zuo), National Science Foundation of China (81770073 to W. Zuo), Shanghai Science and Technology Talents Program (19QB1403100 to W.Zuo), Youth 1000 Talent Plan of China to W. Zuo, Tongji University (Basic Scientific Research Interdisciplinary Fund and 985 Grant to W. Zuo) and Guangzhou Medical University annual grant to W. Zuo. We salute to medical workers who sacrificed their lives in fight against the COVID-19 pandemic.

